# microRNA-19b regulates proliferation & patterning in the avian forebrain

**DOI:** 10.1101/2020.06.21.163980

**Authors:** Archita Mishra, Suvimal Kumar Sindhu, Niveda Udaykumar, Jonaki Sen

## Abstract

Unlike the six-layered organization of neurons in the mammalian neocortex, the avian pallium features spatially-segregated neuronal clusters with partially conserved gene expression. While efforts to uncover similarities between the avian pallium and mammalian neocortex have persisted, the mechanisms of pallial development in birds have not been comprehensively explored. Here, we have established the role of a highly conserved microRNA, miR-19b, in controlling proliferation in the embryonic avian forebrain, by regulating the expression of E2f8, a cell-cycle inhibitor and NeuroD1, a neuronal differentiation factor. Further, investigation revealed that there exists a proliferation-dependent specification of pallial neurons in birds, similar to that in mammals. In summary, these findings suggest a profound role of miR-19b in shaping the avian pallium and indicate a closer resemblance of developmental mechanisms between the avian pallium and the mammalian neocortex.

## Introduction

Elucidating the mechanism(s) driving the evolution of the mammalian laminar neocortex and its relationship to the non-laminar pallium found in birds and reptiles has been a long-standing interest in neuroscience (Butler et al., 2011). The mammalian neocortex features a six-layered laminar pallium, with neurons being born from a common pool of neural stem cells (NSCs)/radial glial cells (RGCs) located in the ventricular zone (VZ). Intermediate basal progenitors (IPCs) produced from the RGCs, reside in the subventricular zone (SVZ), where they undergo a few rounds of cell division to produce more neural progenitor cells (NPCs). These NPCs then differentiate and migrate radially from the dorsal region and tangentially from the ventral region to form neocortical neurons (Haubensak et al., 2004; McConnell & Kaznowski, 1991; Miyata et al., 2004; Molyneaux et al., 2007; Noctor et al., 2004). In contrast, the differentiating immature neurons produced from NPCs in birds, undergo radial migration without the generation of IPCs from RGCs (Cardenas et al., 2018; Nomura et al., 2016). The avian pallium is segregated into clusters of neurons known as nuclei (Ball & Balthazart, 2021; Karten, 1997). Interestingly, in birds laminar organization is observed in limited regions such as the hyperpallium and the auditory cortex (Stacho et al., 2020; Wang et al., 2010). Furthermore, neurons in the developing avian pallium exhibit neocortex-like subtypes, as revealed by marker and projection analyses (Abellan et al., 2014; Atoji & Wild, 2012; Dugas-Ford et al., 2012; Gupta et al., 2012; Puelles et al., 2000; Suzuki & Hirata, 2014; Suzuki et al., 2012). However, the mechanisms governing the production and patterning of these homologous neurons in the avian brain remain poorly understood.

In mammals, several signaling pathways (Ohtaka-Maruyama & Okado, 2015) and microRNAs (miRNAs), which cause post-transcriptional silencing of gene expression of multiple target mRNAs (Agarwal et al., 2015; Bartel, 2004), regulate the production and specification of neocortical neurons (Diaz et al., 2020; Shu et al., 2019; Todorov et al., 2024). Loss of miRNA activity through Dicer ablation reduces the size of the neocortex and negatively impacts NSC survival and differentiation (Andersson et al., 2010; De Pietri Tonelli et al., 2008; Kawase-Koga et al., 2010; Nowakowski et al., 2011). A highly conserved microRNA cluster, miR-17∼92, has been shown to be crucial for survival of mouse embryos (Ventura et al., 2008) and plays a role in expanding NSCs/RGCs in the mammalian brain (Bian et al., 2013). Furthermore, a member of this cluster, miR-19a, regulates the number of NSCs in mammals by controlling the expression of Pten, a tumor suppressor that regulates NSC renewal (Bian et al., 2013; Zheng et al., 2008).

Another member of this cluster miR-19b, an ortholog of miR-19a, has been identified as a prime oncogenic miRNA (Fan et al., 2014; Mendell, 2008; Olive et al., 2009). Our previous study demonstrated that miR-19b is expressed in the VZ of the developing chicken forebrain (Sindhu et al., 2019). These findings motivated us to further investigate the function of miR-19b in regulating neurogenesis. We first conducted a comprehensive expression analysis of miR-19b across different developmental stages in chicken. Interestingly, this analysis revealed an expression pattern that is spatially and temporally synchronous with the neurogenic wave in the developing chicken pallium. Further analysis showed that miR-19b regulates the number of NSCs and controls the expression of E2f8, a crucial regulator of the cell-cycle (Christensen et al., 2005; Maiti et al., 2005), as well as NeuroD1, a key factor in neuronal differentiation (Lee et al., 1995; Pataskar et al., 2016). In fact, we found that altering proliferation and neurogenic differentiation through the functional manipulation of miR-19b affected neuronal patterning in the developing avian forebrain. In summary, this study reveals some conserved mechanisms of neurogenesis and patterning between the avian pallium and the mammalian neocortex during development, reinforcing the notion of their shared evolutionary origin.

## Results

### Mutually exclusive expression of NeuroD1 and miR-19b

We have previously reported that miR-19b is expressed in the VZ of the chick forebrain at Hamilton-Hamburger stage 24 (HH24) (Hamburger & Hamilton, 1951; Sindhu et al., 2019), further analysis revealed that its expression persists in the VZ across various developmental stages e.g. at HH29 and, HH35 (Fig. 1 A-C). Interestingly, NeuroD1 - a transcription factor reported to be necessary and sufficient for neural specification and differentiation, (Lee et al., 1995; Pataskar et al., 2016) is expressed in the subventricular zone (SVZ) in the developing chicken forebrain, in a mutually exclusive expression domain to that of miR-19b, particularly at HH24 (Fig. 1A). Further, at HH29 these two factors are expressed differentially in complementary regions, with miR-19b being expressed at higher levels in the dorsal ventricular ridge (DVR) region (yellow-dashed box) as compared to the wulst region (magenta-dashed box) (Fig. 1B). Interestingly, at HH35, miR-19b expression is relatively low in the wulst region, where NeuroD1 expression is higher and distributed throughout (white arrows in magenta-dashed boxed region) (Fig. 1C). In contrast, NeuroD1 is only present in the SVZ in the DVR region, while miR-19b is expressed at higher levels throughout the DVR region (Fig. 1C). This suggests that miR-19b and NeuroD1 are expressed in a spatio-temporally dynamic manner in mutually exclusive domains in the wulst and DVR region.

**Figure 1.**
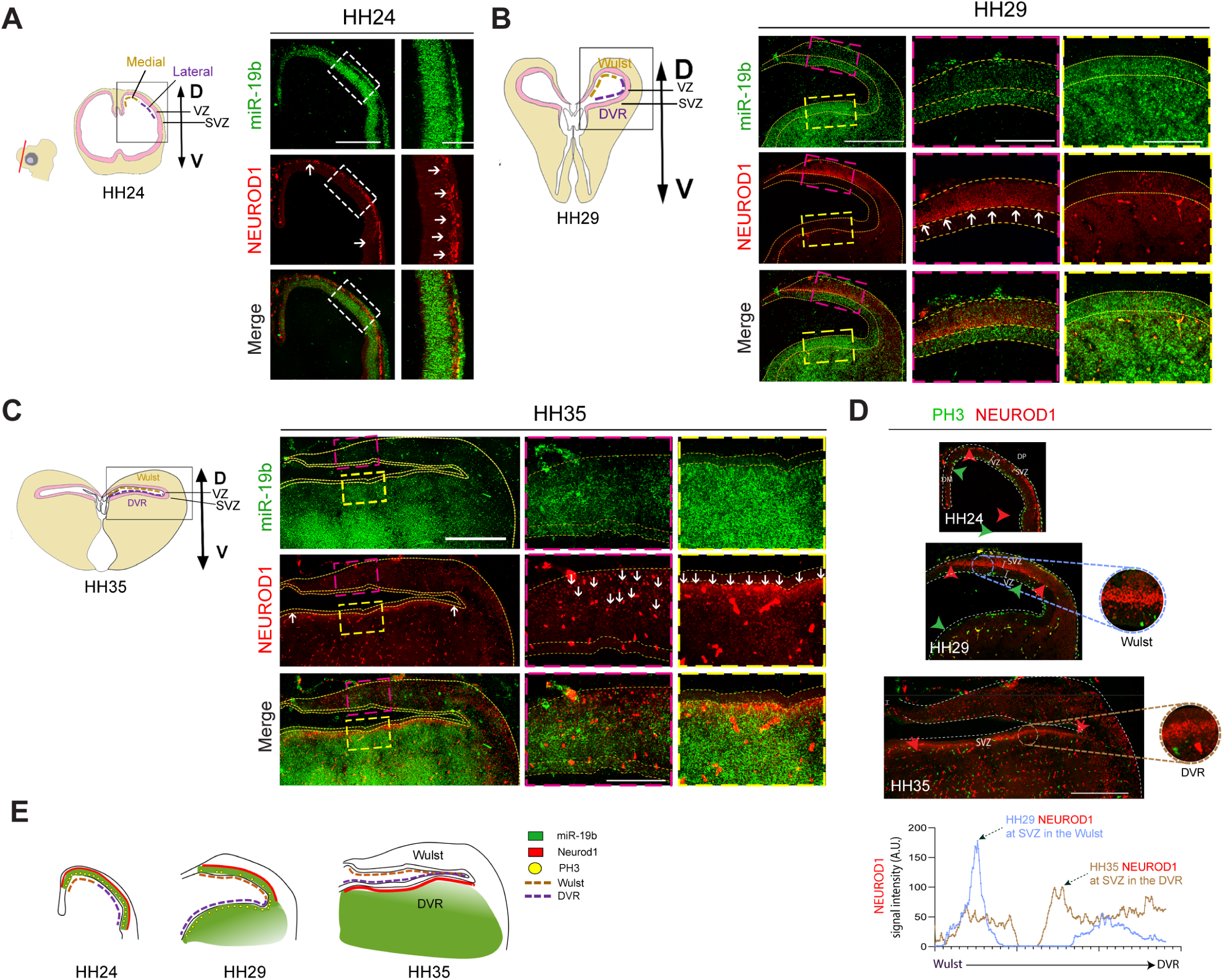
Expression patterns of NEUROD1 and miR-19b in the developing chick pallium and their spatial distribution relative to PH3-positive cells. (A) Screening of NEUROD1 (IHC) and miR-19b (RNA-ISH) expression within chicken pallium at different stages (A) A cross-section of HH24 chicken pallium showing NeuroD1 and miR-19b with a merged image in the inset. Scale bar-500µm, Inset (100µm), n=3 (B) miR-19b and NeuroD1 expression dynamics in wulst (Red inset) and DVR (Yellow inset) region at stage HH29 chicken pallium. Scale bar-500µm, Inset (250µm), n=3 (C) a similar comparison of miR-19b and NeuroD1 expression at stage HH29 chicken pallium. Scale bar - 800µm, inset (250µm), n=3 (D) Distribution of PH3 and NeuroD1-positive cells at all above-mentioned stages domain of PH3 positive cells is marked by green arrows wherein NeuroD1 is represented by red arrows. The intensity plot shows mutually exclusive expression domains of NeuroD1 and miR-19b. Insets depict magnified images. n=3, Scale bar: 800µm. White arrowheads show NeuroD1 positive cells in SVZ. A.U.= arbitrary unit; VZ= ventricular zone, SVZ= sub-ventricular zone.

### Spatio-temporal dynamics of neurogenesis

Neurogenesis in the avian pallium follows a distinct spatio-temporally dynamic pattern, where it begins in the presumptive wulst region followed by the DVR region as is evident from differential labelling with EdU and immunohistochemistry for phospho-Histone 3 (PH3) (Abdel-Mannan et al., 2008; Cheung et al., 2007; Gupta et al., 2012; Suzuki et al., 2012). To correlate this dynamic spatio-temporal pattern of neurogenesis with the complementary expression patterns of miR-19b and NeuroD1, we compared the expression pattern of NeuroD1 with PH3 immunostaining and found that indeed PH3 and NeuroD1 are present in complementary regions across developmental stages. At HH24, PH3+ and NeuroD1+ cells are homogeneously distributed in the VZ and SVZ, marking the presence of proliferating NSCs and post-mitotic immature neurons, respectively (Fig. 1D). However, at HH29 proliferation ceases in the wulst region as demonstrated by low PH3 immunoreactivity and there is a higher expression of NeuroD1 in the SVZ relative to the DVR (Fig. 1D). Further, at HH35 there are negligible number of PH3+ cells in both the regions, whereas DVR shows enriched expression of NeuroD1 in the SVZ. This indicates that while NSC proliferation ceases at both locations, the DVR is still producing post-mitotic immature neurons (Fig. 1D). These findings suggest that the VZ of the presumptive wulst region ceases proliferation earlier than the DVR and NeuroD1 is primarily expressed in SVZ (D’Amico et al., 2013; Lee et al., 2000) of the regions where proliferation have ceased and neuronal differentiation is initiated (FIG. 1E).

Taken together, it appears that the VZ, a proliferative zone, expresses miR-19b whereas the SVZ, a post-mitotic zone, expresses NeuroD1, hence they are in mutually exclusive domains. Hence, we hypothesized that miR-19b might serve as a key regulator of cell proliferation by enhancing proliferation through its expression in the VZ and/or by inhibiting neuronal differentiation through its regulation of NeuroD1, as observed in the mouse pancreas.

### miR-19b as a proliferation regulator

To explore the function(s) of miR-19b, we identified its potential target mRNAs using tools like - miRDB (Chen & Wang, 2020) and TargetScan (Agarwal et al., 2015) and selected 423 candidates from their intersection (Fig. 2A). Gene ontology analysis indicated that miR-19b targets are mostly associated with cellular senescence (Fig. 2B). Enriched expression of miR-19b in the VZ which contains the NSCs, prompted us to explore the possible function of miR-19b in regulation of cell proliferation. Furthermore, among the candidate regulators of cellular-senescence, we chose a cell-cycle inhibitor, E2f8 (Christensen et al., 2005; Maiti et al., 2005), due to its strong expression in the developing chick forebrain at HH30 (Fig. 2E) and previous reports of its regulation by miR-19b in CD4+ T-cells (Lim et al., 2024). TargetScan analysis showed that chicken E2f8 has an evolutionarily conserved target region (Fig. 2 D) (Agarwal et al., 2015). We then performed a sensor assay (Sindhu et al. 2019) to determine if miR-19b regulates the E2f8 transcript via its target regions by co-expressing miR-19b with transcripts of E2f8 fused with an mCherry reporter in HEK293 T cells (Fig. 2C). This resulted in decreased mCherry expression in the presence of miR-19b as compared to the control, suggesting that miR-19b can regulate these transcripts through their target regions *in vitro* (Fig. 2C). Further, *in ovo* gain-of-function (GoF) and loss-of-function (LoF) experiments were performed with overexpression of miR-19b and Sponge-19b, respectively (Sindhu et al., 2019). A Sponge is an artificial transcript that comprises of multiple copies of the miRNA target sequence, which sequester the endogenously expressed functional miRNAs, thus it competitively prevents miRNAs from interacting with its cognate target mRNAs (Ebert & Sharp, 2010). Expression of Sponge-19b that, sequesters miR-19b leading to a decrease in availability of miR-19b in the cell (Sindhu et al., 2019), showed increase in E2f8 transcripts in the developing chick forebrain at HH25 (Fig. 2F). Such manipulations in the levels of miR-19b demonstrated that miR-19b is both necessary as well as sufficient to regulate the expression of E2f8 (Fig. 2E, F). To further establish the role of miR-19b in regulation of the cell-cycle, we employed flow-cytometry to sort chicken fibroblast cells (DF1) and observed that miR-19b over-expression increased the proportion of cells in the S-phase and M-phase, while decreasing the proportion in G0/G1. On the other hand, depletion of functional miR-19b through expression of Sponge-19b had the opposite effect (Fig. 2G). We also performed EdU-labeling of the chick forebrain neurons under conditions of miR-19b GoF and LoF, adding EdU 2 hours before harvesting at HH25. Quantification of GFP+;EdU+ cells showed that EdU incorporation was promoted by miR-19b overexpression, while it was reduced by expression of Sponge-19b (Fig. 2H). In addition, immunohistochemistry for PH3 conducted at HH34 after functional manipulation of the miR-19b, yielded similar results with over-expression of miR-19b increasing GFP+;PH3+ cells, and expression of Sponge-19b leading to a reduction of the same (Fig. 2I). These findings strongly indicate that miR-19b regulates cell proliferation in the developing chick pallium.

**Figure 2.**
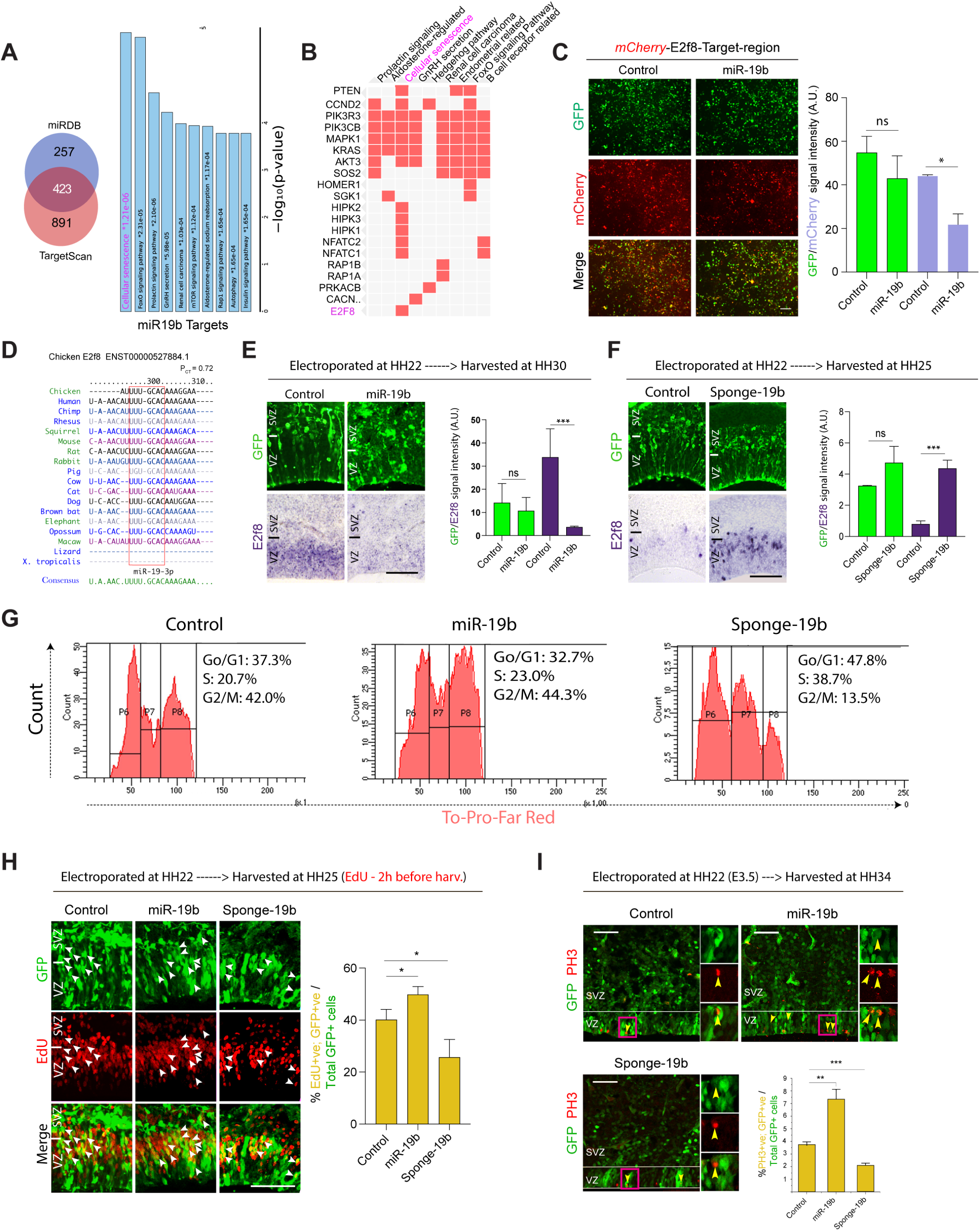
miR-19b regulates E2f8 expression and cell-cycle progression. (A) Venn diagram showing miR-19b targets predicted by miRDB and TargetScan in chicken (*Gallus gallus*) and gene ontology analysis for predicted targets. (B) List of functionally validated targets. The red square indicates the role in the mentioned process/pathway. Potential targets and associated processes highlighted in magenta. (C) Sensor-assay for in vitro demonstration of miR-19b binding to its target region of E2f8. Respective 3’UTRs fused with the coding sequence of mCherry were subjected to miR-19b manipulation. mCherry intensity was further quantified and plotted. Scale bar: 100µm, Plot bar: mean ± s.e.m., N=3 wells, t-test performed, **p≤0.01 and ****p≤0.0001. A.U.= arbitrary unit. (D) represent the conservation of mi19b binding sequences in E2f8 3’ UTR across all species. Effect of miR-19b manipulation on E2f8 (RNA-ISH) (E) and (F), scale bar-300µm. (G) Flow cytometry experiment to analyze cell-cycle distribution of DF1 cells upon manipulation of miR-19b. Go/G1, S and G2/M are presented in population group named as P6, P7 and P8, respectively. Representative images of EdU labeling assay(H) and PH3 immunohistochemistry (I)to demonstrate effect of miR-19b manipulation on cell proliferation. Scale bar: 200µm (H), 50µm (I). n=3, Plot bar: mean ± s.e.m.; A.U.= arbitrary unit. ns= not significant; VZ= ventricular zone, SVZ= sub-ventricular zone. t-test was performed, *p≤0.1, **p≤0.01 and ***p≤0.001, ns = non-significant.

### Perturbation in proliferation affects pallial patterning

Since birds show differential proliferation in two distinct regions of the avian pallium i.e. the wulst and the DVR regions (Abdel-Mannan et al., 2008; Cheung et al., 2007; Gupta et al., 2012; Suzuki et al., 2012) that express separate neocortical layer markers (Dugas-Ford et al., 2012; Puelles et al., 2000; Suzuki & Hirata, 2014; Suzuki et al., 2012), therefore, we wanted to study the relationship between proliferation and specification of distinct regions in the avian pallium. Prior to manipulating proliferation to examine its effect on regional specification, we confirmed the expression of patterning markers - Fezf2 and Mef2c, expressed in the wulst and DVR regions respectively in the avian pallium, which correspond to the deep-layer and upper-layer neurons in the mammalian neocortex (Chen et al., 2008; Molyneaux et al., 2007; Suzuki et al., 2012). Upon analysis, we found that at HH28, Fezf2 was expressed in the ventricular zone (VZ) of the dorsomedial region and the subventricular zone (SVZ) of the dorsal pallium (Fig. 3A). Immunostaining for NeuroD1 - a postmitotic immature neuron marker, confirmed Fezf2+ cells in the SVZ as postmitotic immature neurons, due to their co-expression of NeuroD1 (Fig. 3A, Inset). Additionally, Mef2c expression was observed in cells atop the Fezf2+ cells in the SVZ (Fig. 3A). Thus, Fezf2 and Mef2c expression at HH28 appeared in a bi-laminar form, resembling the neocortical laminar order present in mammals (Molyneaux et al., 2007), indicating the possibility of primitive pallial lamina formation in the common ancestors of birds and mammals. However, at advanced stages such as HH32, HH36 and post-hatching day 0, the expression domains of these markers became spatially segregated, with Fezf2 in the hyperpallium and parahippocampal region (APH) in the wulst and Mef2c in the mesopallium (Msp) in the DVR region (Fig. 3B) (Dugas-Ford et al., 2012; Suzuki et al., 2012). Therefore, as the development of the avian pallium progresses, Fezf2+ and Mef2c+ neurons segregate spatially, departing from their earlier bi-layered arrangement and residing in distinct regions such as presumptive wulst and DVR regions.

**Figure 3.**
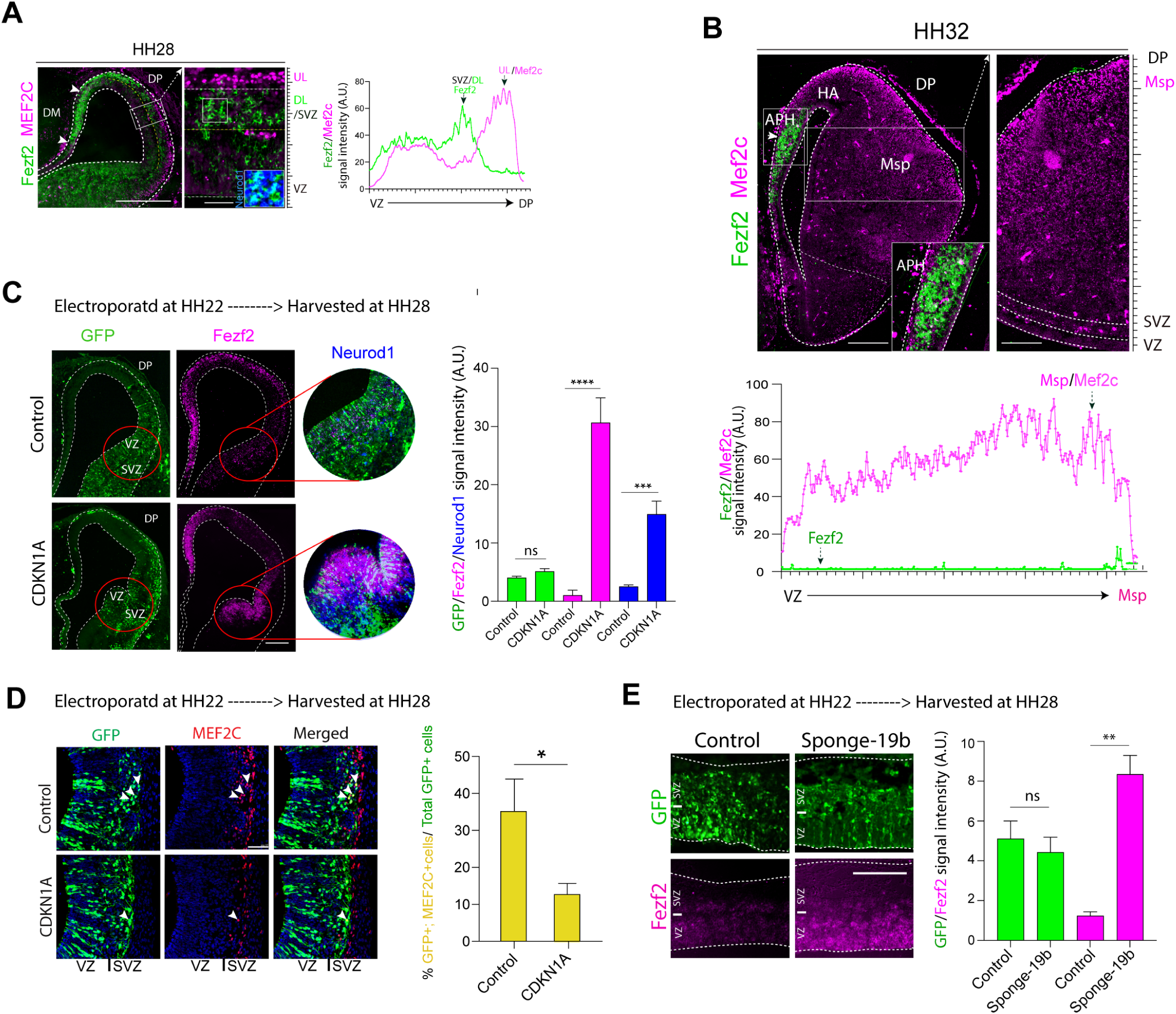
Expression Patterns of Fezf2 and Mef2c in the Embryonic Chick Pallium and the Effects of CDKN1A Overexpression and miR-19b Loss of Function. (A) Expression patterns of Fezf2 and NeuroD1 in the SVZ and Mef2c in the dorsal pallium layer at HH28. The inset highlights double-positive cells for Fezf2 and NeuroD1. Scale bar = 300 µm; 25 µm (inset). n = 3. (B) Fezf2 expression is restricted to the APH, while Mef2c expression extends into the mesopallium (Msp). The inset shows magnified views, and the intensity plot depicts spatial expression profiles of Fezf2 and Mef2c. Scale bar=150µm; n=3; DP= Dorsal Pallium. (C) Effect of ectopic CDKN1A expression on Fezf2(ISH) and NeuroD1(IHC) expression. Electroporation of CDKN1A-overexpressing and control constructs demonstrates altered expression of Fezf2. Magnified regions highlight colocalization of Fezf2 and NeuroD1 (circled regions). Bar graphs display signal intensity quantification; n = 4. Scale bar: 300µm (D) Impact of CDKN1A overexpression on Mef2c expression (IHC). White arrows indicate double-positive cells for Mef2c, and bar graphs quantify their abundance. n=3. (E) Loss of miR-19b affects Fezf2 expression. Signal intensity of Fezf2 per unit electroporated region is quantified and plotted. Bar graph plotted for quantification. n=4, Scale bar: 150µm; Plot bar: mean ± s.e.m. A.U.= arbitrary unit. ns= not significant; VZ= ventricular zone, SVZ= sub-ventricular zone; t-test was performed; *p≤0.1, **p≤0.01 and ***p≤0.001; ns = non-significant.

To test the role of proliferation in patterning, we inhibited the cell-proliferation by overexpressing CDKN1A gene that encodes P21 protein, a cell-cycle inhibitor (Deng et al., 1995; Harper et al., 1993; Zhang et al., 1999), and found ectopic expression of Fezf2 and NeuroD1, indicating that inhibiting the cell-cycle alone is sufficient to generate these neuronal subtypes in the developing chick pallium (Fig. 3C). Interestingly, chicken NPCs that exit the cell-cycle earlier than other NPCs *in vitro* have been shown to preferentially produce more Fezf2+ neurons at the expense of Mef2c+ neurons, much like in mammals (Shen et al., 2006; Suzuki et al., 2012). Thus, *in vitro* findings support the *in vivo* evidence that the early cessation of proliferation leads to the production of Fezf2+ neurons. Further analysis of the CDKN1A overexpressing regions revealed a decrease in the Mef2c+ neuronal population (Fig. 3D). These results indicate that precocious cessation of proliferation leads to preferential production of deep-layer like Fezf2+ neurons and suppresses the alternative upper-layer-like Mef2c+ fate (Chen et al., 2008; McConnell & Kaznowski, 1991).

Our observation of cell-cycle inhibition resulting in ectopic Fezf2 expression (Fig. 3C), along with the demonstration of miR-19b as a cell proliferation regulator, prompted us to investigate if miR-19b might also regulate the expression of deep-layer neuronal markers. We conducted a LoF experiment for miR-19b by expressing Sponge-19b in the VZ and interestingly this resulted in ectopic Fezf2 expression in comparison to the control (Fig. 3E). This suggests that miR-19b plays a crucial role in specification of deep-layer-like neurons in the chick pallium. Such mechanism of proliferation-dependent specification of pallial neurons resembles what is observed in the mammalian neocortex (Shen et al., 2006) suggesting conservation in the mechanisms for cortical neuronal specification between birds and mammals.

### miR-19b regulates NeuroD1 which in turn regulates pallial patterning

Interestingly, miR-19b has been shown to also regulate the NeuroD1 transcript, albeit in the mouse pancreas (Zhang et al., 2011). Based on this and the mutually exclusive expression pattern of miR-19b and NeuroD1 (Fig. 1A-E, 4A), along with the fact that chicken NeuroD1 mRNAs have evolutionarily conserved miR-19b binding sites (Fig. 4B), we hypothesized that miR-19b regulates NeuroD1 in the developing chicken forebrain. A sensor assay for NeuroD1 was performed, similar to the one described for E2f8 (Fig. 2C) and we found that indeed miR-19b can regulate the NeuroD1-target region in a sequence-dependent manner *in vitro* (Fig. 4 C). However, experiments carried out *in ovo* suggested that while miR-19b is sufficient for regulating the expression of NeuroD1, it is not necessary for the same (Fig. 4D-E). This is because miR-19b over-expression could decrease NeuroD1+ cells whereas Sponge-19b expression did not lead to an increase in NeuroD1+ cells relative to the control (Fig. 4D-E). This may be explained by the presence of other inhibitors of NeuroD1 *in vivo*.

**Figure 4.**
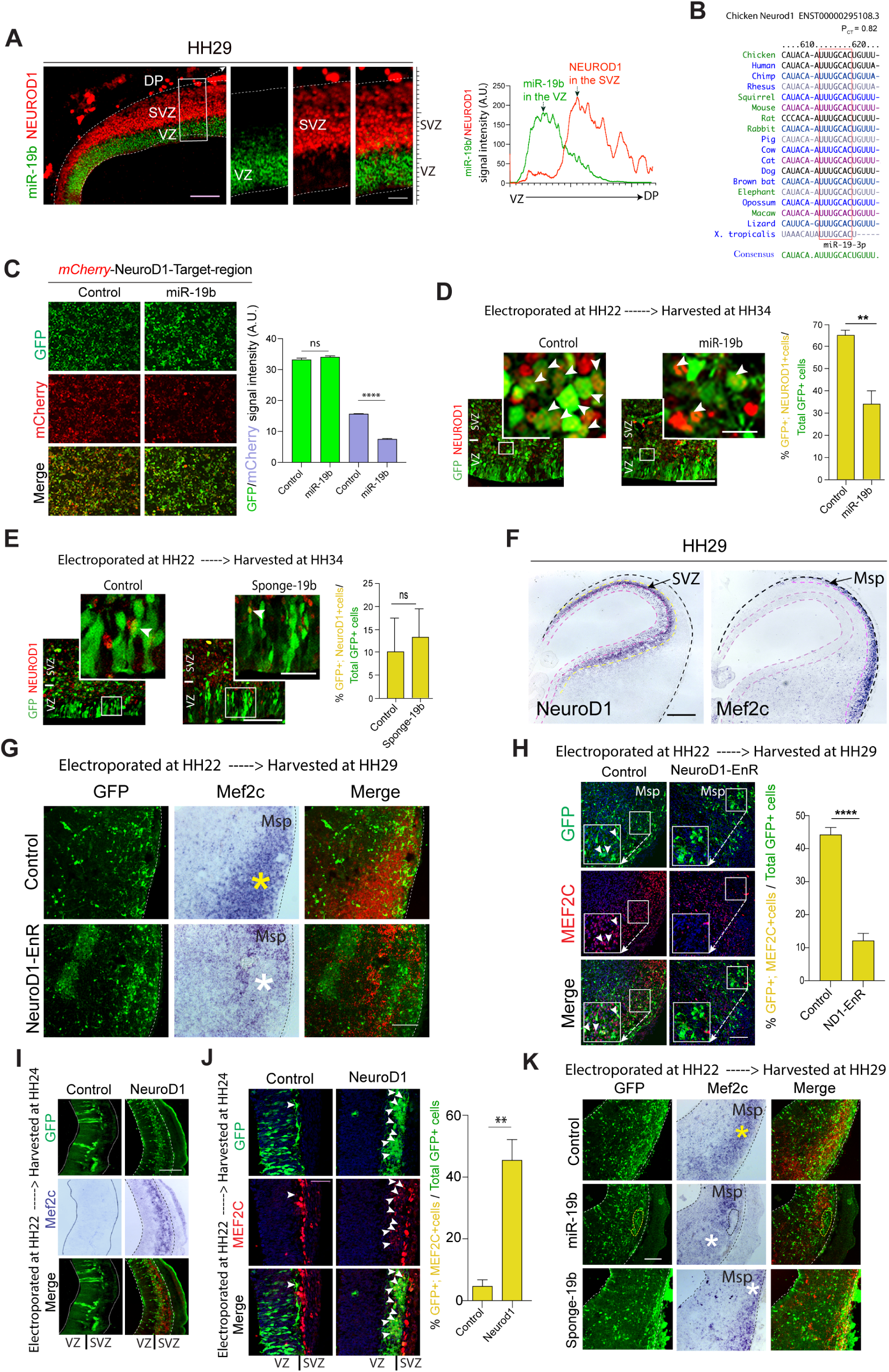
miR-19b Mediated Regulation of NeuroD1 and Its Impact on Patterning in the Developing Chick Pallium. (A) mRNA in situ hybridization (ISH) screening of miR-19b and NeuroD1 immunostaining in the chick pallium at HH29. The intensity plot highlights mutually exclusive expression domains of NeuroD1 and miR-19b. Insets show magnified views of the expression patterns. Scale bar = 100µm; 30µm (insets), n = 3. (B) Conservation analysis of miR-19b binding sequences in the NeuroD1 3′ UTR across species. (C) Sensor assay demonstrating in vitro binding of miR-19b to the target region of NeuroD1. The 3′ UTR of NeuroD1 was fused with mCherry and subjected to miR-19b manipulation. mCherry intensity was quantified and plotted. Scale bar = 100 µm, with n = 3 wells. A.U. = arbitrary units. (D, E) miR-19b regulates NeuroD1 expression in *in vivo* as shown by immunostaining in gain-of-function (D) and loss-of-function (E) experiments. Scale bar = 100 µm, n=4. (F) Expression patterns of NeuroD1 and Mef2c in the chick pallium at HH29. Scale bar = 500µm. (G, H) NeuroD1 loss-of-function experiment using NeuroD1-DBD-EnR electroporation shows loss of Mef2c ISH expression. (G) and IHC expression (H). The yellow asterisk (control) and white asterisk (test) mark the Mef2c mRNA expression domains. Scale bar:100µm, n=3 (I-J) NeuroD1 gain-of-function led to ectopic Mef2c mRNA (I) and Proteins (J) shown by ISH and immunostaining (arrowheads show ectopic expression of Mef2c). n=3, scale bar: 100µm. (K) Effect of miR-19b overexpression and Sponge-19b expression on Mef2c expression. Yellow asterisk is showing Mef2c ISH in control and white asterisk show decreased Mef2c expression using ISH; n=3; Scale bar:100µm; Plot bar: mean ± s.e.m. A.U.= arbitrary unit. ns= not significant; VZ= ventricular zone, SVZ= sub-ventricular zone; Msp = Mesopallium; t-test was performed; *p≤0.1, **p≤0.01 and ***p≤0.001; ns = non-significant.

Since NeuroD1 has also been implicated in causing cell-cycle exit and promoting neural specification and differentiation (Lee et al., 1995; Pataskar et al., 2016) and cell-cycle perturbation affects avian pallial patterning (Fig. 3C-D), therefore we wanted to find out if manipulation of NeuroD1 expression can also affect patterning in the avian pallium. Prior to this, we established the relationship of expression domains of lateral pallial marker i.e. Mef2c with NeuroD1 to better score the changes in regional expression of Mef2c upon perturbation of NeuroD1. The expression pattern comparison of NeuroD1 and Mef2c in the forebrain at HH29 showed that NeuroD1 is expressed in the SVZ region, while Mef2c is expressed in the mesopallium region (Fig. 4F). This suggests that NeuroD1 is transiently expressed in immature neurons, followed by Mef2c as they mature, similar to the upper layer of the mouse neocortex (Lee et al., 1995; Molyneaux et al., 2007; Pataskar et al., 2016). Subsequently, to test the role of NeuroD1 in patterning, we manipulated the function of NeuroD1 by expressing a dominant-negative version, NeuroD1-EnR, where DNA-binding domain of chicken NeuroD1 was fused to the Engrailed-Repressor domain (Yan & Wang, 2004) and found that this indeed inhibits the expression of Mef2c mRNA as well as protein (Fig. 4G, H). On the other hand, NeuroD1 overexpression induced the expression of Mef2c mRNA and Protein (Fig. 4I). These data indicate that NeuroD1 is both necessary and sufficient to regulate the expression of Mef2c, which is crucial for pallial patterning.

Since miR-19b is sufficient to regulate NeuroD1 expression (Fig. 4D), which in turn regulates Mef2c expression (Fig. 4G-J), we directly tested the effect of manipulating miR-19b on Mef2c expression. We observed that upon miR-19b GoF there was a down-regulation of Mef2c expression while miR-19b LoF, mediated by expression of Sponge-19b, led to a decrease in Mef2c expression (Fig. 4K). The ability of Sponge-19b to decrease Mef2C was surprising as we had shown earlier that it does not downregulate NeuroD1 (Fig 4E). The possible explanation for this could be as follows: Sponge-19b can inhibit proliferation (Fig. 2H-I) and proliferation inhibition leads to induction of Fezf2, while inhibiting Mef2c expression (Fig. 3C-D). Thus, it is likely that the decrease in Mef2c expression with Sponge-19b could result from inhibition of cell proliferation rather than through inhibition of NeuroD1 expression.

## Discussion

The avian pallial organization and the mammalian neocortical arrangement are quite different, despite sharing homologous neurons. Therefore, in this study, we examined the chicken pallium to explore key mechanisms of neurogenesis and patterning that may be evolutionarily conserved and hence influence mammalian neocortical development. Here, we have uncovered the following striking similarities between the two species, in terms of mechanisms of production of neurons: a) miR-19a/b-mediated control of neurogenic proliferation, b) miR-19b-mediated repression of NeuroD1 expression and c) proliferation-dependent specification of neurons leading to regulation of pallial patterning. In addition, we have also found a transient pallial bi-layer comprising of Fezf2+ and Mef2c+ neurons in the early embryonic chick forebrain. Overall, this study provides strong evidence that miR-19b could be a major player in shaping avian pallial development using developmental mechanisms partially conserved in mammals.

### miR-19b regulates proliferation of neural progenitor cells

This study has revealed that miR-19b is expressed in the VZ and has been shown to regulate the NSCs number. Further, it also targets E2f8, an inhibitor of cell-proliferation and NeuroD1, a neuronal differentiation factor. Thus, miR-19b is very likely to be a crucial regulator of proliferation in the developing chicken forebrain.

The E2f family of transcription factors regulates the cell-cycle, with some acting as activators and others as repressors (Maiti et al., 2005). Atypical member of E2f family, E2f8, along with E2f7, supports cell-survival in developing mouse embryos (Li et al., 2008) and inhibits proliferation of skin cancer cells (Thurlings et al., 2017). E2f8 has been shown to act as inhibitor of transcription as well (Park et al., 2022). Additionally, miR-19b has been also shown to repress expression of E2f8 in CD4+ T-cell (Lim et al., 2024).Therefore, E2f8 is likely to play a similar role in the chick forebrain, in regulating cell proliferation by promoting cell-cycle exit, and its control by miR-19b may significantly impact neuron production in the avian embryonic pallium.

NeuroD1 has been shown to initiate the neuronal differentiation program (Lee et al., 1995; Pataskar et al., 2016; Schwab et al., 2000) and promote cell-cycle exit, when misexpressed in the mitotic retinal cells (Ochocinska & Hitchcock, 2009). Therefore, suppressing NeuroD1 in the VZ may be essential for maintaining the proliferative state of NSCs, and the role of miR-19b could be important in this context.

Based on our observation that miR-19b inhibits E2f8 expression as well as NeuroD1 expression, we speculate that miR-19b is required for maintaining the proliferative state of NPCs by blocking their cell-cycle exit through suppression of E2f8 and preventing premature neurogenesis through inhibition of NeuroD1 expression in the VZ. All the above support our hypothesis that miR-19b regulates cell proliferation in the VZ in the developing chick forebrain. Moreover, since the ortholog of miR-19b i.e. miR-19a has also been shown to regulate neurogenesis (Bian et al., 2013), our study suggests that miR19-mediated mechanism of regulation of neurogenesis is conserved between birds and mammals.

### Progression of the cell-cycle and specification of neocortical neurons

In the mouse there is an intrinsic program that regulates sequential formation of deep-layer and upper-layer neurons in a temporally regulated manner (Shen et al., 2006). A similar process has been reported for *in vitro* neurogenesis from chicken pallial neural stem cell (Suzuki et al., 2012).

Overexpression of CyclinD1-Cdk4 in the mouse embryonic pallium increases Tbr2+ basal progenitors (IPCs) in the SVZ, initially inhibiting neuronal differentiation (Lange et al., 2009). Over time, CyclinD1-Cdk4 effects weaken, leading to more IPCs and mature upper-layer neurons in the cortical plate. Ectopic expression of CyclinD1-Cdk4 in chick embryos at HH24 also produces postmitotic cells that are Tbr2+ immature neurons, although their subtypes were not further identified to explore the link between cell-progression and neuronal specification (Nomura et al., 2016).

In our study, we found that in the early chick embryo suppressing the progression of cell-cycle led to the production of postmitotic neurons, as shown by NeuroD1 expression along with Fezf2 in these cells. This is similar to what has been observed when chick NSCs at an early stage produce deep-layer-like Fezf2+ neurons under *in vitro* conditions (Suzuki et al., 2012). Interestingly, in mice, the gene CDKN1B, which is related to CDKN1A, produces the P27 protein, a cell-cycle inhibitor. P27 is important for the formation of upper-layer neurons because it slows down the cell-cycle and helps activate Ngn2, which controls the expression of NeuroD1 (Nguyen et al., 2006; Roybon et al., 2009). Thus, there is a striking similarity between the chick and the mouse with respect to the link between the time of exit from cell-cycle and the specification of neuronal subtype.

### miR19b as a regulator of NeuroD1 expression

NeuroD1 has been previously shown to be both necessary and sufficient for neural specification and differentiation (Lee et al., 1995; Pataskar et al., 2016). Recently, it has been demonstrated that transient expression of NeuroD1 is sufficient to activate the neurogenic program in progenitor cells (Pataskar et al., 2016). Therefore, it is likely that one or more factors in the neural progenitor cells in the VZ maybe necessary to inhibit any precocious expression of NeuroD1.

Our study suggests that miR-19b is sufficient but not necessary to regulate NeuroD1 expression. This could possibly be due to the presence of other inhibitory factors such as other microRNAs and transcriptional repressors, in the VZ. In mammals, additional microRNAs, such as miR30a-5p (Kim et al., 2013), miR-190 (Zheng et al., 2010) and miR-124 (Liu et al., 2011), have been reported to regulate the expression of NeuroD1 in various tissue contexts. Hence, it is possible that one or more of these microRNAs are present in the VZ of the developing chick forebrain and contribute to the inhibition of NeuroD1 expression, both under natural conditions and in experimental settings involving the suppression of miR-19b.

### NeuroD1 and production of neocortical neurons

NeuroD1 was discovered as a gene that is both necessary and sufficient for the differentiation of neurons in several regions of the nervous system, including the forebrain (Boutin et al., 2010; Gao et al., 2009; Lee et al., 1995; Schwab et al., 2000). Interestingly, NeuroD1 GoF experiments in the mouse brain showed increased cell-cycle exit and more postmitotic neurons in the intermediate zones compared to controls, although layer markers were not analyzed separately (Pataskar et al., 2016). However, NeuroD1 overexpression in the muller cells in retina showed its re-specification into different neuronal subtypes depending on its expression level (Xu et al., 2023). In our study, we found that NeuroD1 is both necessary and sufficient for the expression of Mef2c+ upper-layer-like neocortical neurons in the chick pallium. Hence, NeuroD1 appears to play a key role in regulating production and specification of neurons during pallial development.

In conclusion, our study establishes miR-19b as a significant contributor to the regulation of NSC proliferation, differentiation, and pallial patterning in birds in an evolutionarily conserved manner. Thus, our findings provide novel insight into some aspects of shared evolutionary and developmental mechanisms that regulate pallial development.

## Materials and methods

### Key resource table

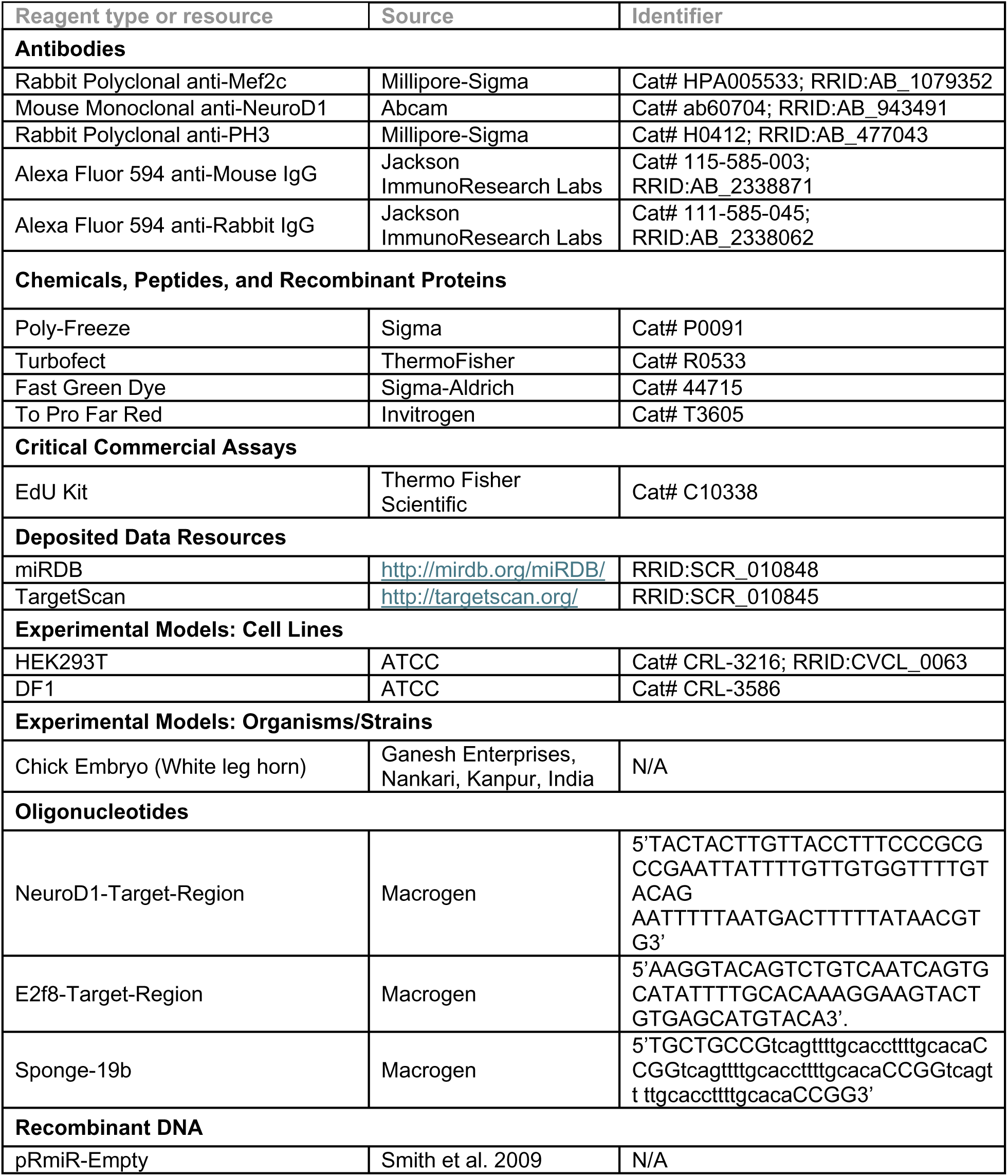
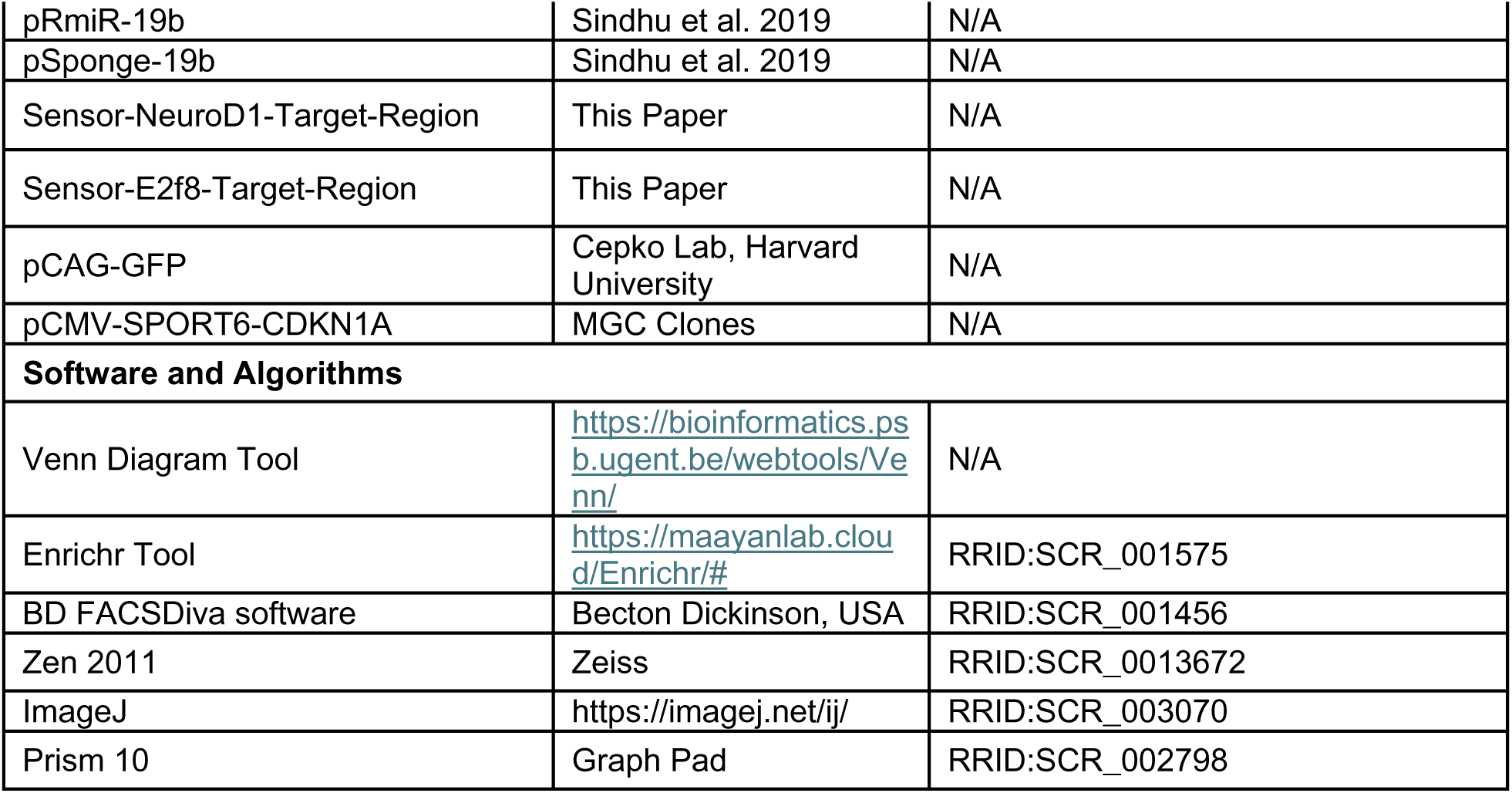

### Animals

Fertilized chicken eggs from the White Leghorn variety were purchased from Ganesh Enterprises, Nankari, Indian Institute of Technology Kanpur, Uttar Pradesh, India. These eggs were incubated in a humidified incubator at 38 °C until desired Hamilton Hamburger (HH) stages (Hamburger and Hamilton 1992).

### Cell-culture

Human embryonic kidney (HEK-293T) (ATCC; CRL-3216) and chicken embryonic fibroblast, DF-1 (ATCC; CRL-3586) cells were cultured with 10% Fetal Bovine Serum in Dulbecco’s Modified Eagle’s Medium (DMEM) media.

### Tissue preparation

The chick embryo at the desired HH stages were taken and brains were dissected out. The fixation was performed at 4°C overnight in 4% Paraformaldehyde (PFA). These brains were washed with phosphate buffer saline (PBS), and after dehydration was performed in 30% sucrose, they were embedded in Poly-freeze (Sigma, P0091). Finally, a cryostat (Leica CM 1850) was used to obtain 12-16 μm thick sections of the chick pallium.

### RNA *in situ* hybridization

For microRNA *in situ* hybridization, a probe was designed where the 3’end of the antisense sequence of miR-19b was labeled with digoxigenin (Eurofins MWG Operon). This probe was used to perform section RNA *in situ* hybridization using a published protocol (Thompson, Deo, and Turner 2007). Finally, the detection was performed using another published method (Trimarchi et al. 2007).

The mRNA *in situ* hybridization was performed on the tissue sections using a published method (Trimarchi et al. 2007). The following cDNA clones were used to prepare probes: (1) Fezf2 (pTA2-cFezf2) was a gift from Prof. T. Hirata, National Insitutute of Genetics, Japan, (2) E2f8 (ChEST 638e24) was procured from Chicken EST collection from Source Biosciences (previously MRC Geneservice, UK).

### *In silico* analysis of miR-19b target mRNAs

The list of target mRNAs was obtained using Targetscan & miRDB. Using venn diagram tool (https://bioinformatics.psb.ugent.be/webtools/Venn/) intersection revealed 423 target mRNAs. After that, plot & gene ontology analysis were performed using online tool Enrichr (https://maayanlab.cloud/Enrichr/<u>#</u>).

### Plasmid construction

1. pRmiR-19b: pRmiR-Empty construct (Smith et al. 2009) after BsaI digestion, was ligated with the synthesized primary sequence of miR-19b (Macrogen Inc.) to generate pRmiR-19b.
2. Sensor-ND1-Target-Region: The chicken NeuroD1 3’UTR having miR-19b target region with flanking NotI and HindIII sites were synthesized (Macrogen Inc.) and ligated in pCAG-mCherry construct (a gift from Prof. Constance Cepko, Harvard Medical School, USA), downstream of the mCherry. Chicken NeuroD1 target region: (NM_204920.1 from 1380^th^ nucleotide position to 1458^th^ nucleotide position) 5’TACTACTTGTTACCTTTCCCGCGCCGAATTATTTTGTTGTGGTTTTGTACAGAATTTTTAATGACTTTTTATAACGTG3’.
3. Sensor-E2f8-Target-Region: The chicken E2f8 3’UTR having miR-19b target region with flanking NotI and HindIII sites were synthesized (Macrogen Inc.) and ligated in pCAG-mCherry construct (a gift from Prof. Constance Cepko, Harvard Medical School, USA), downstream of the mCherry. Chicken E2f8 target region: XM_040672655.2 from 3108^th^ nucleotide position to 3167^th^ nucleotide position) 5’AAGGTACAGTCTGTCAATCAGTGCATATTTTGCACAAAGGAAGTACTGTGAGCATGTACA3’.
4. pSponge-19b: For this first three copies of miR-19b target sequence (written in small letters) was ligated in the pRmiR vector (Smith et al. 2009) to obtain pSponge-19b-3x construct 5’TGCTGCCGtcagttttgcaccttttgcacaCCGGtcagttttgcaccttttgcacaCCGGtcagtt ttgcaccttttgcacaCCGG3’. After that, using SalI, BglII, and BamHI Sponge-19b-3X sequence was chained in tandem to obtain twelve copies of sponge-19b and this construct was named pSponge-19b (Ebert and Sharp 2010) results in loss-of-function of miR-19b.
5. pCAG-GFP was a gift from Prof. C. Cepko, Harvard Medical School, USA.
6. pCMV-SPORT6-CDKN1A from MGC clones.

### Sensor assay

HEK-293T cells were transfected using Turbofect (Thermo Fisher Scientific Inc., R0533) as per manufacturer’s protocol, with pRmiR vector(s) and respective Sensor vector(s) following which GFP and mCherry fluorescence were observed after 72 hours.

### *In ovo* electroporation

Fertilized chicken eggs were incubated overnight and lowered by taking out 2-3ml of albumin. Then, a window cut in the shell on the top and into one of the forebrain vesicles the DNA construct (2μg/μl) with 0.1% of fast green (Sigma-Aldrich, 44715) was injected. Platinum hockey stick electrodes (Nepagene, Japan) were used to deliver an electric current of 13V in five pulses for 50ms duration with a gap of 950ms, to the HH22 chick embryo head, using an electroporator (ECM830 Harvard instruments, USA). Before sealing with the tape, Penicillin, and Streptomycin with sterile PBS, were added over the embryos. Finally, harvesting was done at the desired HH stage.

### Immunofluorescence

Briefly, after 4% PFA fixation for 5 minutes, three PBT washes were performed. Blocking was performed with 5% of heat-inactivated goat serum (HINGS) for 1 hour. Primary antibody was added and incubated overnight at 4°C. The primary antibodies were used at the following dilution: anti-Mef2c, 1:200 (Millipore-Sigma, HPA005533); anti-NeuroD1, 1:1500 (Abcam, ab60704), anti-PH3, 1:300 (Sigma, H0412). After washing three times in PBT for five minutes each, the appropriate secondary antibody was added. Secondary antibodies used were as follows: Alexa Flour 594-conjugated Anti-Mouse IgG, 1:250 (Jackson ImmunoResearch Laboratories, INC., 115-585-003) and Alexa Flour 594-conjugated Anti-Rabbit IgG, 1:250 (Jackson ImmunoResearch Laboratories, INC., 111-585-045).

### Cell-Cycle analysis

Chicken embryonic fibroblast, DF1 cells were transfected with respective DNA constructs using Turbofect (Thermo Fisher Scientific Inc., R0533), as per manufacturer’s protocol. Cells were harvested after Trypsin treatment for 2-4 minutes then after cells were washed with DMEM media. After that cells were passed through strainer to obtain single-cell suspension, followed by treatment with 70% ice-cold ethanol for 30 minutes at 4dC. After that, two washes of PBT (PBS with 0.02% Tween-20) were done. After that cells were treated with RNase A (10mg/ml) for 30 minutes at room temperature. For labelling 10μl of 10/ng/μl To-Pro-Far Red (Invitrogen, T3605) was added to the cells for 15 minutes at room temperature after that washed with PBS. Fluorescence of individual cells were quantified using flow cytometer (BD FACS Calibur, Becton Dickinson, USA) using the facility at Central Drug Research Institute, Lucknow, India. Data were analyzed using BD FACSDiva software (Becton Dickinson, USA).

### EdU labeling and detection

After electroporation of desired DNA constructs at HH23, and then 2 hours before harvesting 25μg of EdU (ThermoFisher Scientific, C10338) was overlaid in 200μl of 1X Phosphate Buffer Saline (PBS). After harvesting at E4.5 (HH25), brains were dissected out in PBS and emersion fixed in 4% paraformaldehyde and after sequential dehydration with up to 30% sucrose, brains were embedded in the Poly-freeze media (Millipore-Sigma, SHH0026) and 14 micrometer sections were generated and mounted on glass slide. EdU detection was performed on these sections using CLICK-IT EdU-labeling kit (ThermoFisher Scientific, C10338) as per manufacturer’s protocol.

### Image acquisition and statistical analysis

To obtain the images of RNA *in situ* hybridization as well as immunofluorescence a Leica stereomicroscope (DM500B) equipped with a DFC500 camera was used. Other fluorescent images were obtained using the multi-photon laser scanning confocal microscope LSM780 (Carl Zeiss Inc.) and ZEN 2011 software.

Each set of experiments was repeated at least three times; for animals, n=1 means on one animal experiment was performed; for cell culture, n=1 means in one dish experiment was performed. Only samples with GFP and/or mCherry expression after transfection or electroporation were included in the analysis. ImageJ software was used to measure the fluorescence intensity. Graph Pad Prism 10 were used to plot the graph and to perform unpaired student’s t-test and calculate the significance.

## Acknowledgements

We acknowledge Prof. Amitabha Bandhopadhyay for suggestions on the manuscript and Ms. Neetu Dey (IIT Kanpur, India) for microscopy. Mr. D.N. Vishwakarma (CDRI, Lucknow, India) for flow-cytometry. Funded by SERB-DST, Govt. of India. (EMR/2016/000886) to J.S. CSIR, Govt. of India to S.K.S. for PhD (09/092(0824)/2011-EMR-I). MHRD, Govt. of India. to A.M. and N.U. for PhD.

## Additional information

### Funding

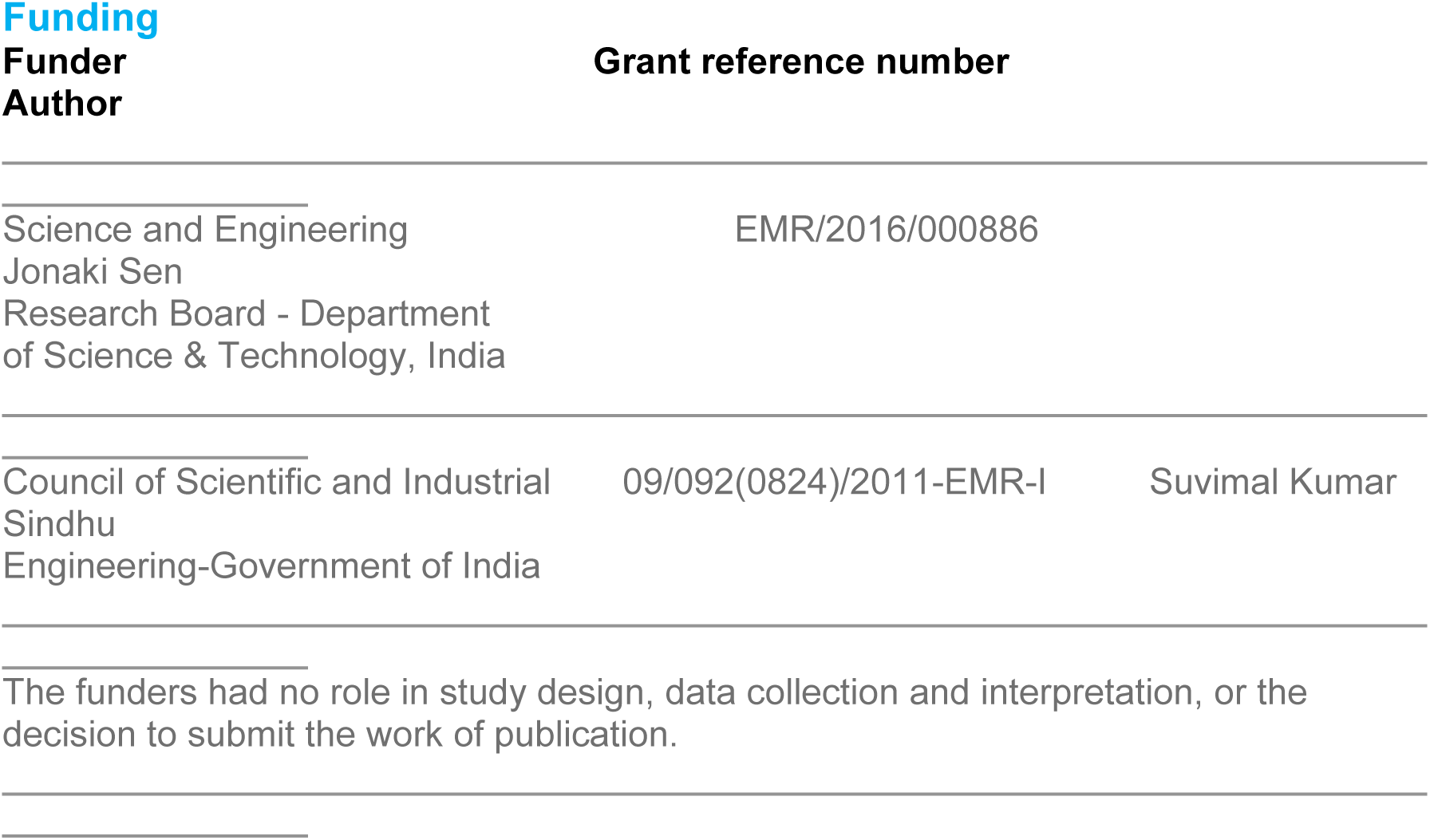

### Author contributions

S.K.S., and J.S. designed study; S.K.S., A.M., N.U. performed experiments; S.K.S., A.M., N.U., and J.S. analyzed data; J.S. Supervised; S.K.S., A.M., and J.S. wrote the paper.

### Ethics

All experiments conducted in this study were in accordance with the guidelines of Indian Institute of Technology Kanpur, India.

## Additional files

### Data availability

All data generated or analyzed during this study are including in the manuscript and supporting files.

## References

Abdel-Mannan, O., Cheung, A. F., & Molnar, Z. (2008). Evolution of cortical neurogenesis. Brain Res Bull, 75(2-4), 398–404. 10.1016/j.brainresbull.2007.10.047

Abellan, A., Desfilis, E., & Medina, L. (2014). Combinatorial expression of Lef1, Lhx2, Lhx5, Lhx9, Lmo3, Lmo4, and Prox1 helps to identify comparable subdivisions in the developing hippocampal formation of mouse and chicken. Front Neuroanat, 8, 59. 10.3389/fnana.2014.00059

Agarwal, V., Bell, G. W., Nam, J. W., & Bartel, D. P. (2015). Predicting effective microRNA target sites in mammalian mRNAs. Elife, 4. 10.7554/eLife.05005

Andersson, T., Rahman, S., Sansom, S. N., Alsio, J. M., Kaneda, M., Smith, J., O’Carroll, D., Tarakhovsky, A., & Livesey, F. J. (2010). Reversible block of mouse neural stem cell differentiation in the absence of dicer and microRNAs. PLoS One, 5(10), e13453. 10.1371/journal.pone.0013453

Atoji, Y., & Wild, J. M. (2012). Afferent and efferent projections of the mesopallium in the pigeon (Columba livia). J Comp Neurol, 520(4), 717–741. 10.1002/cne.22763

Ball, G. F., & Balthazart, J. (2021). Evolutionary neuroscience: Are the brains of birds and mammals really so different? Curr Biol, 31(13), R840–R842. 10.1016/j.cub.2021.05.004

Bartel, D. P. (2004). MicroRNAs: genomics, biogenesis, mechanism, and function. Cell, 116(2), 281–297. 10.1016/s0092-8674(04)00045-5

Bian, S., Hong, J., Li, Q., Schebelle, L., Pollock, A., Knauss, J. L., Garg, V., & Sun, T. (2013). MicroRNA cluster miR-17-92 regulates neural stem cell expansion and transition to intermediate progenitors in the developing mouse neocortex. Cell Rep, 3(5), 1398–1406. 10.1016/j.celrep.2013.03.037

Boutin, C., Hardt, O., de Chevigny, A., Core, N., Goebbels, S., Seidenfaden, R., Bosio, A., & Cremer, H. (2010). NeuroD1 induces terminal neuronal differentiation in olfactory neurogenesis. Proc Natl Acad Sci U S A, 107(3), 1201–1206. 10.1073/pnas.0909015107

Butler, A. B., Reiner, A., & Karten, H. J. (2011). Evolution of the amniote pallium and the origins of mammalian neocortex. Ann N Y Acad Sci, 1225, 14–27. 10.1111/j.1749-6632.2011.06006.x

Cardenas, A., Villalba, A., de Juan Romero, C., Pico, E., Kyrousi, C., Tzika, A. C., Tessier-Lavigne, M., Ma, L., Drukker, M., Cappello, S., & Borrell, V. (2018). Evolution of Cortical Neurogenesis in Amniotes Controlled by Robo Signaling Levels. Cell, 174(3), 590–606 e521. 10.1016/j.cell.2018.06.007

Chen, B., Wang, S. S., Hattox, A. M., Rayburn, H., Nelson, S. B., & McConnell, S. K. (2008). The Fezf2-Ctip2 genetic pathway regulates the fate choice of subcortical projection neurons in the developing cerebral cortex. Proc Natl Acad Sci U S A, 105(32), 11382–11387. 10.1073/pnas.0804918105

Chen, Y., & Wang, X. (2020). miRDB: an online database for prediction of functional microRNA targets. Nucleic Acids Res, 48(D1), D127–D131. 10.1093/nar/gkz757

Cheung, A. F., Pollen, A. A., Tavare, A., DeProto, J., & Molnar, Z. (2007). Comparative aspects of cortical neurogenesis in vertebrates. J Anat, 211(2), 164–176. 10.1111/j.1469-7580.2007.00769.x

Christensen, J., Cloos, P., Toftegaard, U., Klinkenberg, D., Bracken, A. P., Trinh, E., Heeran, M., Di Stefano, L., & Helin, K. (2005). Characterization of E2F8, a novel E2F-like cell-cycle regulated repressor of E2F-activated transcription. Nucleic Acids Res, 33(17), 5458–5470. 10.1093/nar/gki855

D’Amico, L. A., Boujard, D., & Coumailleau, P. (2013). The neurogenic factor NeuroD1 is expressed in post-mitotic cells during juvenile and adult Xenopus neurogenesis and not in progenitor or radial glial cells. PLoS One, 8(6), e66487. 10.1371/journal.pone.0066487

De Pietri Tonelli, D., Pulvers, J. N., Haffner, C., Murchison, E. P., Hannon, G. J., & Huttner, W. B. (2008). miRNAs are essential for survival and differentiation of newborn neurons but not for expansion of neural progenitors during early neurogenesis in the mouse embryonic neocortex. Development, 135(23), 3911–3921. 10.1242/dev.025080

Deng, C., Zhang, P., Harper, J. W., Elledge, S. J., & Leder, P. (1995). Mice lacking p21CIP1/WAF1 undergo normal development, but are defective in G1 checkpoint control. Cell, 82(4), 675–684. 10.1016/0092-8674(95)90039-x

Diaz, J. L., Siththanandan, V. B., Lu, V., Gonzalez-Nava, N., Pasquina, L., MacDonald, J. L., Woodworth, M. B., Ozkan, A., Nair, R., He, Z., Sahni, V., Sarnow, P., Palmer, T. D., Macklis, J. D., & Tharin, S. (2020). An evolutionarily acquired microRNA shapes development of mammalian cortical projections. Proc Natl Acad Sci U S A, 117(46), 29113–29122. 10.1073/pnas.2006700117

Dugas-Ford, J., Rowell, J. J., & Ragsdale, C. W. (2012). Cell-type homologies and the origins of the neocortex. Proc Natl Acad Sci U S A, 109(42), 16974–16979. 10.1073/pnas.1204773109

Ebert, M. S., & Sharp, P. A. (2010). MicroRNA sponges: progress and possibilities. RNA, 16(11), 2043–2050. 10.1261/rna.2414110

Fan, Y., Yin, S., Hao, Y., Yang, J., Zhang, H., Sun, C., Ma, M., Chang, Q., & Xi, J. J. (2014). miR-19b promotes tumor growth and metastasis via targeting TP53. RNA, 20(6), 765–772. 10.1261/rna.043026.113

Gao, Z., Ure, K., Ables, J. L., Lagace, D. C., Nave, K. A., Goebbels, S., Eisch, A. J., & Hsieh, J. (2009). Neurod1 is essential for the survival and maturation of adult-born neurons. Nat Neurosci, 12(9), 1090–1092. 10.1038/nn.2385

Gupta, S., Maurya, R., Saxena, M., & Sen, J. (2012). Defining structural homology between the mammalian and avian hippocampus through conserved gene expression patterns observed in the chick embryo. Dev Biol, 366(2), 125–141. 10.1016/j.ydbio.2012.03.027

Hamburger, V., & Hamilton, H. L. (1951). A series of normal stages in the development of the chick embryo. J Morphol, 88(1), 49–92. https://www.ncbi.nlm.nih.gov/pubmed/24539719

Harper, J. W., Adami, G. R., Wei, N., Keyomarsi, K., & Elledge, S. J. (1993). The p21 Cdk-interacting protein Cip1 is a potent inhibitor of G1 cyclin-dependent kinases. Cell, 75(4), 805–816. 10.1016/0092-8674(93)90499-g

Haubensak, W., Attardo, A., Denk, W., & Huttner, W. B. (2004). Neurons arise in the basal neuroepithelium of the early mammalian telencephalon: a major site of neurogenesis. Proc Natl Acad Sci U S A, 101(9), 3196–3201. 10.1073/pnas.0308600100

Karten, H. J. (1997). Evolutionary developmental biology meets the brain: the origins of mammalian cortex. Proc Natl Acad Sci U S A, 94(7), 2800–2804. 10.1073/pnas.94.7.2800

Kawase-Koga, Y., Low, R., Otaegi, G., Pollock, A., Deng, H., Eisenhaber, F., Maurer-Stroh, S., & Sun, T. (2010). RNAase-III enzyme Dicer maintains signaling pathways for differentiation and survival in mouse cortical neural stem cells. J Cell Sci, 123(Pt 4), 586–594. 10.1242/jcs.059659

Kim, J. W., You, Y. H., Jung, S., Suh-Kim, H., Lee, I. K., Cho, J. H., & Yoon, K. H. (2013). miRNA-30a-5p-mediated silencing of Beta2/NeuroD expression is an important initial event of glucotoxicity-induced beta cell dysfunction in rodent models. Diabetologia, 56(4), 847–855. 10.1007/s00125-012-2812-x

Lange, C., Huttner, W. B., & Calegari, F. (2009). Cdk4/cyclinD1 overexpression in neural stem cells shortens G1, delays neurogenesis, and promotes the generation and expansion of basal progenitors. Cell Stem Cell, 5(3), 320–331. 10.1016/j.stem.2009.05.026

Lee, J. E., Hollenberg, S. M., Snider, L., Turner, D. L., Lipnick, N., & Weintraub, H. (1995). Conversion of Xenopus ectoderm into neurons by NeuroD, a basic helix-loop-helix protein. Science, 268(5212), 836–844. 10.1126/science.7754368

Lee, J. K., Cho, J. H., Hwang, W. S., Lee, Y. D., Reu, D. S., & Suh-Kim, H. (2000). Expression of neuroD/BETA2 in mitotic and postmitotic neuronal cells during the development of nervous system. Dev Dyn, 217(4), 361–367. 10.1002/(SICI)1097-0177(200004)217:4<361::AID-DVDY3>3.0.CO;2-8

Li, J., Ran, C., Li, E., Gordon, F., Comstock, G., Siddiqui, H., Cleghorn, W., Chen, H. Z., Kornacker, K., Liu, C. G., Pandit, S. K., Khanizadeh, M., Weinstein, M., Leone, G., & de Bruin, A. (2008). Synergistic function of E2F7 and E2F8 is essential for cell survival and embryonic development. Dev Cell, 14(1), 62–75. 10.1016/j.devcel.2007.10.017

Lim, Y. J., Park, S. A., Wang, D., Jin, W., Ku, W. L., Zhang, D., Xu, J., Patino, L. C., Liu, N., Chen, W., Kazmi, R., Zhao, K., Zhang, Y. E., Sun, L., & Chen, W. (2024). MicroRNA-19b exacerbates systemic sclerosis through promoting Th9 cells. Cell Rep, 43(8), 114565. 10.1016/j.celrep.2024.114565

Liu, K., Liu, Y., Mo, W., Qiu, R., Wang, X., Wu, J. Y., & He, R. (2011). MiR-124 regulates early neurogenesis in the optic vesicle and forebrain, targeting NeuroD1. Nucleic Acids Res, 39(7), 2869–2879. 10.1093/nar/gkq904

Maiti, B., Li, J., de Bruin, A., Gordon, F., Timmers, C., Opavsky, R., Patil, K., Tuttle, J., Cleghorn, W., & Leone, G. (2005). Cloning and characterization of mouse E2F8, a novel mammalian E2F family member capable of blocking cellular proliferation. J Biol Chem, 280(18), 18211–18220. 10.1074/jbc.M501410200

McConnell, S. K., & Kaznowski, C. E. (1991). Cell cycle dependence of laminar determination in developing neocortex. Science, 254(5029), 282–285. 10.1126/science.254.5029.282

Mendell, J. T. (2008). miRiad roles for the miR-17-92 cluster in development and disease. Cell, 133(2), 217–222. 10.1016/j.cell.2008.04.001

Miyata, T., Kawaguchi, A., Saito, K., Kawano, M., Muto, T., & Ogawa, M. (2004). Asymmetric production of surface-dividing and non-surface-dividing cortical progenitor cells. Development, 131(13), 3133–3145. 10.1242/dev.01173

Molyneaux, B. J., Arlotta, P., Menezes, J. R., & Macklis, J. D. (2007). Neuronal subtype specification in the cerebral cortex. Nat Rev Neurosci, 8(6), 427–437. 10.1038/nrn2151

Nguyen, L., Besson, A., Heng, J. I., Schuurmans, C., Teboul, L., Parras, C., Philpott, A., Roberts, J. M., & Guillemot, F. (2006). p27kip1 independently promotes neuronal differentiation and migration in the cerebral cortex. Genes Dev, 20(11), 1511–1524. 10.1101/gad.377106

Noctor, S. C., Martinez-Cerdeno, V., Ivic, L., & Kriegstein, A. R. (2004). Cortical neurons arise in symmetric and asymmetric division zones and migrate through specific phases. Nat Neurosci, 7(2), 136–144. 10.1038/nn1172

Nomura, T., Ohtaka-Maruyama, C., Yamashita, W., Wakamatsu, Y., Murakami, Y., Calegari, F., Suzuki, K., Gotoh, H., & Ono, K. (2016). The evolution of basal progenitors in the developing non-mammalian brain. Development, 143(1), 66–74. 10.1242/dev.127100

Nowakowski, T. J., Mysiak, K. S., Pratt, T., & Price, D. J. (2011). Functional dicer is necessary for appropriate specification of radial glia during early development of mouse telencephalon. PLoS One, 6(8), e23013. 10.1371/journal.pone.0023013

Ochocinska, M. J., & Hitchcock, P. F. (2009). NeuroD regulates proliferation of photoreceptor progenitors in the retina of the zebrafish. Mech Dev, 126(3-4), 128–141. 10.1016/j.mod.2008.11.009

Ohtaka-Maruyama, C., & Okado, H. (2015). Molecular Pathways Underlying Projection Neuron Production and Migration during Cerebral Cortical Development. Front Neurosci, 9, 447. 10.3389/fnins.2015.00447

Olive, V., Bennett, M. J., Walker, J. C., Ma, C., Jiang, I., Cordon-Cardo, C., Li, Q. J., Lowe, S. W., Hannon, G. J., & He, L. (2009). miR-19 is a key oncogenic component of mir-17-92. Genes Dev, 23(24), 2839–2849. 10.1101/gad.1861409

Park, S. A., Lim, Y. J., Ku, W. L., Zhang, D., Cui, K., Tang, L. Y., Chia, C., Zanvit, P., Chen, Z., Jin, W., Wang, D., Xu, J., Liu, O., Wang, F., Cain, A., Guo, N., Nakatsukasa, H., Wu, C., Zhang, Y. E.,…Chen, W. (2022). Opposing functions of circadian protein DBP and atypical E2F family E2F8 in anti-tumor Th9 cell differentiation. Nat Commun, 13(1), 6069. 10.1038/s41467-022-33733-8

Pataskar, A., Jung, J., Smialowski, P., Noack, F., Calegari, F., Straub, T., & Tiwari, V. K. (2016). NeuroD1 reprograms chromatin and transcription factor landscapes to induce the neuronal program. EMBO J, 35(1), 24–45. 10.15252/embj.201591206

Puelles, L., Kuwana, E., Puelles, E., Bulfone, A., Shimamura, K., Keleher, J., Smiga, S., & Rubenstein, J. L. (2000). Pallial and subpallial derivatives in the embryonic chick and mouse telencephalon, traced by the expression of the genes Dlx-2, Emx-1, Nkx-2.1, Pax-6, and Tbr-1. J Comp Neurol, 424(3), 409–438. 10.1002/1096-9861(20000828)424:3<409::aid-cne3>30.co;2-7

Roybon, L., Hjalt, T., Stott, S., Guillemot, F., Li, J. Y., & Brundin, P. (2009). Neurogenin2 directs granule neuroblast production and amplification while NeuroD1 specifies neuronal fate during hippocampal neurogenesis. PLoS One, 4(3), e4779. 10.1371/journal.pone.0004779

Schwab, M. H., Bartholomae, A., Heimrich, B., Feldmeyer, D., Druffel-Augustin, S., Goebbels, S., Naya, F. J., Zhao, S., Frotscher, M., Tsai, M. J., & Nave, K. A. (2000). Neuronal basic helix-loop-helix proteins (NEX and BETA2/Neuro D) regulate terminal granule cell differentiation in the hippocampus. J Neurosci, 20(10), 3714–3724. 10.1523/JNEUROSCI.20-10-03714.2000

Shen, Q., Wang, Y., Dimos, J. T., Fasano, C. A., Phoenix, T. N., Lemischka, I. R., Ivanova, N. B., Stifani, S., Morrisey, E. E., & Temple, S. (2006). The timing of cortical neurogenesis is encoded within lineages of individual progenitor cells. Nat Neurosci, 9(6), 743–751. 10.1038/nn1694

Shu, P., Wu, C., Ruan, X., Liu, W., Hou, L., Fu, H., Wang, M., Liu, C., Zeng, Y., Chen, P., Yin, B., Yuan, J., Qiang, B., Peng, X., & Zhong, W. (2019). Opposing Gradients of MicroRNA Expression Temporally Pattern Layer Formation in the Developing Neocortex. Dev Cell, 49(5), 764–785 e764. 10.1016/j.devcel.2019.04.017

Sindhu, S. K., Udaykumar, N., Zaidi, M. A. A., Soni, A., & Sen, J. (2019). MicroRNA-19b restricts Wnt7b to the hem, which induces aspects of hippocampus development in the avian forebrain. Development, 146(20). 10.1242/dev.175729

Stacho, M., Herold, C., Rook, N., Wagner, H., Axer, M., Amunts, K., & Gunturkun, O. (2020). A cortex-like canonical circuit in the avian forebrain. Science, 369(6511). 10.1126/science.abc5534

Suzuki, I. K., & Hirata, T. (2014). A common developmental plan for neocortical gene-expressing neurons in the pallium of the domestic chicken Gallus gallus domesticus and the Chinese softshell turtle Pelodiscus sinensis. Front Neuroanat, 8, 20. 10.3389/fnana.2014.00020

Suzuki, I. K., Kawasaki, T., Gojobori, T., & Hirata, T. (2012). The temporal sequence of the mammalian neocortical neurogenetic program drives mediolateral pattern in the chick pallium. Dev Cell, 22(4), 863–870. 10.1016/j.devcel.2012.01.004

Thurlings, I., Martinez-Lopez, L. M., Westendorp, B., Zijp, M., Kuiper, R., Tooten, P., Kent, L. N., Leone, G., Vos, H. J., Burgering, B., & de Bruin, A. (2017). Synergistic functions of E2F7 and E2F8 are critical to suppress stress-induced skin cancer. Oncogene, 36(6), 829–839. 10.1038/onc.2016.251

Todorov, H., Weissbach, S., Schlichtholz, L., Mueller, H., Hartwich, D., Gerber, S., & Winter, J. (2024). Stage-specific expression patterns and co-targeting relationships among miRNAs in the developing mouse cerebral cortex. Commun Biol, 7(1), 1366. 10.1038/s42003-024-07092-7

Ventura, A., Young, A. G., Winslow, M. M., Lintault, L., Meissner, A., Erkeland, S. J., Newman, J., Bronson, R. T., Crowley, D., Stone, J. R., Jaenisch, R., Sharp, P. A., & Jacks, T. (2008). Targeted deletion reveals essential and overlapping functions of the miR-17 through 92 family of miRNA clusters. Cell, 132(5), 875–886. 10.1016/j.cell.2008.02.019

Wang, Y., Brzozowska-Prechtl, A., & Karten, H. J. (2010). Laminar and columnar auditory cortex in avian brain. Proc Natl Acad Sci U S A, 107(28), 12676–12681. 10.1073/pnas.1006645107

Xu, D., Zhong, L. T., Cheng, H. Y., Wang, Z. Q., Chen, X. M., Feng, A. Y., Chen, W. Y., Chen, G., & Xu, Y. (2023). Overexpressing NeuroD1 reprograms Muller cells into various types of retinal neurons. Neural Regen Res, 18(5), 1124–1131. 10.4103/1673-5374.355818

Yan, R. T., & Wang, S. Z. (2004). Requirement of neuroD for photoreceptor formation in the chick retina. Invest Ophthalmol Vis Sci, 45(1), 48–58. 10.1167/iovs.03-0774

Zhang, P., Wong, C., Liu, D., Finegold, M., Harper, J. W., & Elledge, S. J. (1999). p21(CIP1) and p57(KIP2) control muscle differentiation at the myogenin step. Genes Dev, 13(2), 213–224. 10.1101/gad.13.2.213

Zhang, Z. W., Zhang, L. Q., Ding, L., Wang, F., Sun, Y. J., An, Y., Zhao, Y., Li, Y. H., & Teng, C. B. (2011). MicroRNA-19b downregulates insulin 1 through targeting transcription factor NeuroD1. FEBS Lett, 585(16), 2592–2598. 10.1016/j.febslet.2011.06.039

Zheng, H., Ying, H., Yan, H., Kimmelman, A. C., Hiller, D. J., Chen, A. J., Perry, S. R., Tonon, G., Chu, G. C., Ding, Z., Stommel, J. M., Dunn, K. L., Wiedemeyer, R., You, M. J., Brennan, C., Wang, Y. A., Ligon, K. L., Wong, W. H., Chin, L., & DePinho, R. A. (2008). p53 and Pten control neural and glioma stem/progenitor cell renewal and differentiation. Nature, 455(7216), 1129–1133. 10.1038/nature07443

Zheng, H., Zeng, Y., Zhang, X., Chu, J., Loh, H. H., & Law, P. Y. (2010). mu-Opioid receptor agonists differentially regulate the expression of miR-190 and NeuroD. Mol Pharmacol, 77(1), 102–109. 10.1124/mol.109.060848

